# Natural infection with *Giardia* is associated with altered community structure of the human and canine gut microbiome

**DOI:** 10.1101/2020.01.13.905604

**Authors:** Alexander S.F. Berry, Kaylynn Johnson, Rene Martins, Megan Sullivan, Camila Farias Amorim, Alexandra Putre, Aiysha Scott, Shuai Wang, Brianna Lindsay, Robert Baldassano, Thomas J. Nolan, Daniel P. Beiting

## Abstract

Enteric parasitic infections are among the most prevalent infections in lower- and middle-income countries (LMICs), and have a profound impact on global public health. While the microbiome is increasingly recognized as a key determinant of gut health and human development, the impact of naturally-acquired parasite infections on microbial community structure in the gut, and the extent to which parasite-induced changes in the microbiome may contribute to gastrointestinal symptoms, is poorly understood. Enteric parasites are routinely identified in companion animals in the United States, presenting a unique opportunity to leverage this animal model to investigate the impact of naturally-acquired parasite infections on the microbiome. Clinical, parasitological, and microbiome profiling of a cohort of 258 dogs revealed a significant correlation between parasite infection and composition of the bacterial community in the gut. Relative to other enteric pathogens, *Giardia* was associated with a more pronounced perturbation of the microbiome. Using a database mining approach that allowed us to compare our findings to a large-scale epidemiological study of enteric diseases in humans, we also observed a substantial alteration to microbiome structure in *Giardia*-infected children. Importantly, infection was associated with a reduction in the relative abundance of potential pathobionts, including *Gammaproteobacteria*, and an increase in *Prevotella* - a profile often associated with gut health. Taken together, our data show that widespread *Giardia* infection in young animals and humans is associated with significant remodeling of the gut microbiome, and provide a possible explanation for the high prevalence of asymptomatic *Giardia* infections observed across host species.

**Importance:** While enteric parasitic infections are among the most important infections in lower- and middle-income countries, their impact on gut microbiota is poorly understood. We reasoned that clinical symptoms associated with these infections may be influenced by alterations of the microbiome that occur during infection. To explore this notion, we took a two-pronged approach. First, we studied a cohort of dogs naturally infected with various enteric parasites and found a strong association between parasite infection and altered gut microbiota composition. *Giardia*, one of the most prevalent parasite infections globally, had a particularly large impact on the microbiome. Second, we took a database-driven strategy to integrate microbiome data with clinical data from large human field studies and found that *Giardia* infection is also associated with marked alteration of the gut microbiome of children, suggesting a possible explanation for why *Giardia* has been reported to be associated with protection from moderate-to-severe diarrhea.

## Introduction

Enteric parasites, including helminths and protozoa, are among the most prevalent infections in lower- and middle-income countries (LMICs) with an estimated 3.5 billion people affected worldwide (1, 2). Infection with eukaryotic pathogens often results in acute, moderate-to-severe diarrheal disease and/or chronic malnutrition and stunting, which has significant consequences for morbidity and mortality (3–5). Conversely, some intestinal parasites are frequently associated with asymptomatic infections, or are even considered commensal (6, 7). *Giardia*, for example, was found in 18 of 1093 (1.6%) of healthy volunteers in Melbourne, Australia (8) and in 286 of 1359 (21%) of healthy schoolchildren in Madrid, Spain (9). It is important to understand whether and how these abundant and pervasive parasites impact gut health.

While the microbiome is increasingly recognized as a key determinant of gut health and human development, the impact of naturally-acquired parasite infections on the microbial community in the gut is poorly understood. Many studies of parasites and their impact on the microbiome involve experimental infections of laboratory mice (10–12). While such studies can be powerful for elucidating mechanism, they often involve laboratory-adapted parasite strains, specialized animal husbandry practices, or high infectious doses, all of which can impact host immunity and the composition of the microbiome. Conversely, studies of parasite infections in human populations are challenging due to the relatively low prevalence of these infections in developed countries and the presence of confounding variables, such as malnourishment (13–16). These issues are largely overcome by studying enteric parasite infections in companion animals. Various enteric parasites are frequently found in screenings of domestic dogs and cats in the United States (17). For example, a study of over one million dogs throughout the United States in 2006 found that 12.5% were infected with at least one enteric parasite, with the most prevalent being *Giardia* which infected 4% of dogs (18). As companion animals, dogs are increasingly recognized as an ideal model system for translational gut microbiome research. In addition to a harboring similar gut microbiota as humans, dogs often share their environment with humans, consume a similar omnivorous diet, and can spontaneously develop GI disease that shares many features in common with inflammatory bowel disease in humans (19–26). In addition, like humans, dogs frequently become infected with enteric parasites in early life. Here we performed 16S rRNA sequencing of fecal samples from 258 dogs naturally-infected with one or more eukaryotic parasites to evaluate the impact of parasite infection on gut microbiota composition. We found that parasite infections significantly perturb the microbiome and that *Giardia* is associated with the largest changes in canine gut microbiota.

We also investigated whether *Giardia* – a frequent infection among humans residing in LMICs – causes similar perturbations in human gut microbiota composition. The Global Enteric Multicenter Study (GEMS) investigated the causes of pediatric moderate-to-severe diarrhea (MSD) in LMICs (27). In addition to reporting a strong association between infection with rotavirus or *Cryptosporidium* and the development of MSD, this study also described the surprising observation that *Giardia* was significantly associated with protection from MSD. A follow-up study performed 16S sequencing of fecal samples from approximately 1000 GEMS participants (28), but this study only considered the relationship between the microbiome and MSD, and did not examine a role for parasite infections in influencing this relationship. We used a database mining approach to determine if *Giardia* infection perturbs the human gut microbiome similarly to how it perturbs the canine gut microbiome, and to gain insight into possible mechanisms by which *Giardia* infection is linked to protection against diarrhea.

## Results

### Enteric parasite infections perturb the canine microbiome

A stool bank was generated from samples screened at a veterinary clinical parasitology service as part of our Companion Animal Microbiome during Parasitism (CAMP) study (see methods). A total of 258 canine fecal samples were split into 9 groups based on parasite-infection status (**Fig. 1A**): No parasite seen (NPS) controls, *Giardia*, *Cystoisospora*, hookworm, whipworm, ascarid, tapeworm, *Eucoleus boehmi*, and co-infections. Since certain enteric parasites, such as *Giardia*, are more prevalent in young animals, we sought to control for age and other potential confounding variables in our statistical analyses. Parasite infection status is associated with significant changes in beta diversity, as determined by both Bray-Curtis and weighted UniFrac, even when covariates such as age, sex, and spay/neuter status were controlled for as confounding variables (p<0.05, PERMANOVA) (**Fig. 1B**). Approximately 5% of the variation in microbiome composition was explained by parasite infection status, compared to <1% explained by age, sex, or spay/neuter status (p>0.05, PERMANOVA)(**Fig. 1B**). Specifically, *Giardia-* and co-infected animals displayed the most significantly differences in beta diversity compared to NPS controls using both Bray-Curtis (**Fig. 1C**) and weighted UniFrac metrics (**Fig. 1D**).

**Figure 1.**
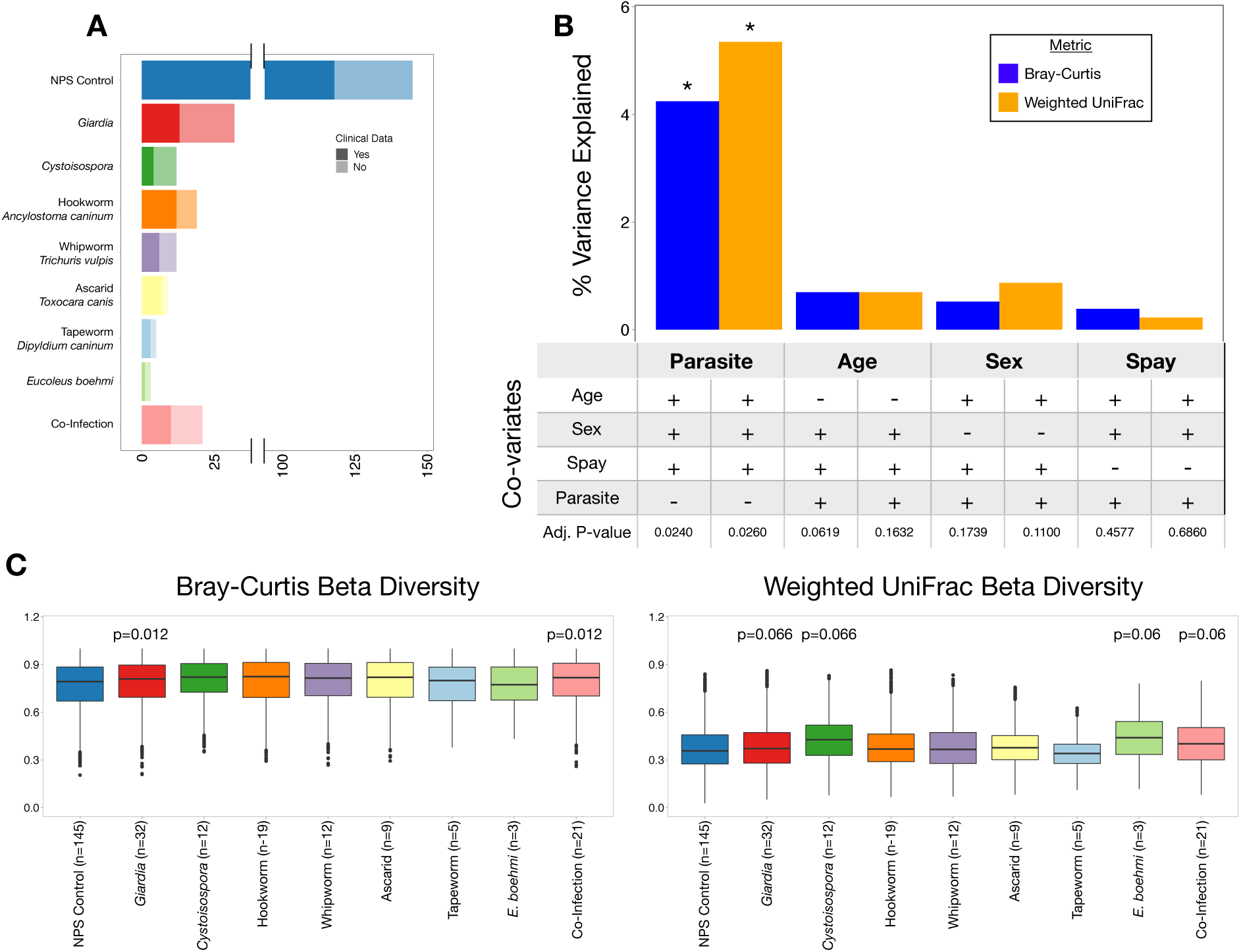
Parasite infection perturbs canine gut microbiota. 16S sequencing of fecal samples from 258 dogs infected with none, one, or multiple enteric parasites was performed. A) The number of no parasite seen (NPS) controls, dogs infected with each parasite, and dogs infected with more than one parasite are shown. The number of dogs for which clinical data is unavailable are greyed out. B) The percent variance in Bray-Curtis and weighted UniFrac beta diversity explained by each variable is represented by blue and orange bars, respectively. Whether or not age, sex, spay/neuter status, or parasite infection were controlled, and the significance of each variable are shown below each bar. Asterisks highlight variables with adjusted p < 0.05. C) Boxplots show the difference in Bray-Curtis beta diversity and D) weighted UniFrac beta diversity between samples in each infection and NPS Controls. Adjusted p-values < 0.1 are labeled above each box.

### Canine *Giardia* infection is associated with significant alterations in gut microbiota composition

Given the diverse range of parasites detected in our animals, we set out to determine whether specific types of parasites were associated with more pronounced microbiome alterations. *Giardia* infection is associated with a change in Bray-Curtis (p<0.01, 1.6% of total variation) and weighted UniFrac (p<0.05, 1.5% of total variation) beta diversity compared to NPS controls, without controlling for age, sex, and spay/neuter status (**Fig. 2A**). When controlling for age, sex, and spay/neuter status, beta diversity is still significantly altered during *Giardia* infection as measured by Bray-Curtis (p<0.05, 1.1% of total variation), but no longer meets the 0.05 cutoff for significance for weighted UniFrac (p=0.0997, 1.0% of total variation) (**Fig. 2A**). The differences in beta diversity between *Giardia* infection and NPS controls were driven by several bacterial taxa as determined by LDA Effect Size (LEfSe) analysis (**Figs. 2B and 2C**). *Giardia* is associated with enrichment of *Clostridium*, a genus that contains several commensal taxa, as well as an enrichment of *Lactobacillus*. However, *Giardia* was also associated with a reduction in *Bacteroides*, a genus that includes important commensal bacteria. In order to verify the taxa associated with *Giardia* infection, point-biserial correlation coefficients were calculated for each taxa with average relative abundance <1%. Consistent with our LEfSe results, point-biserial correlation coefficients also showed enrichment of *Clostridium* and *Lactobacillus*, and a reduction in *Bacteroides* in addition to a reduction in *Megamonas* (**Table S1**). The high relative abundance of *Clostridium* and *Lactobacillus*, and the low relative abundance of *Bacteroides* in *Giardia*-infected dogs compared with NPS controls shows that *Giardia* infection in animals is associated with an altered gut microbiota composition.

**Figure 2.**
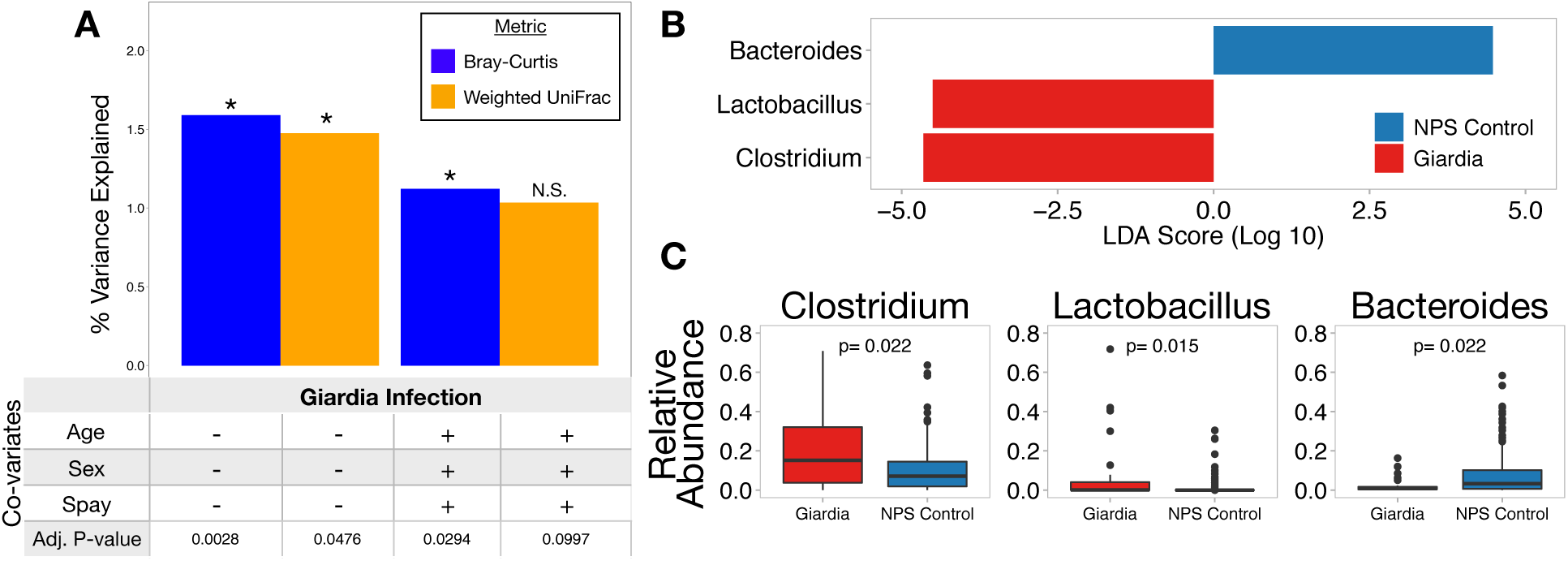
Giardia infection is associated with enrichment of several key bacterial taxa in the canine gut. A) Histogram shows that *Giardia* infection is associated with a significant difference in beta diversity compared to NPS controls. Bar height reflects the percent of total beta diversity variance is explained by *Giardia* infection. Pluses underneath show when age, sex, and spay/neuter status are controlled for. Asterisks denote bars with adjusted p < 0.05. B) LEfSe graph shows the magnitude of enrichment with LDA Score > 2 comparing *Giardia*-infected dogs to NPS control dogs. C) Boxplots show the relative abundance of differentially-enriched taxa. *Clostridium* is among the most highly enriched bacterial taxa associated with *Giardia* infection compared to controls.

It is possible that the microbiota changes observed in *Giardia-*infected dogs could be driven, in part, by clinical variables such as diarrhea or antibiotic use. To discriminate between microbiota changes linked to diarrhea or antibiotics versus those linked to infection, we evaluated medical records, when available (n=174), to identify animals with a recent history of diarrhea or antibiotic use (**Fig. 1A**). Interestingly, *Giardia* infection was not associated with diarrhea or antibiotic use (chi-squared test, p>0.25): among dogs with clinical data, four of 13 (30.1%) *Giardia*-infected dogs had diarrhea compared to 21 of 118 (18%) NPS control dogs with diarrhea (**Fig. 3A**), while two of 13 (15.4%) *Giardia*-infected dogs received antibiotics compared to 15 of 118 (12.7%) NPS control dogs (**Fig S1A**). In contrast, antibiotic use was strongly correlated with diarrhea (chi-squared test, p<0.01), with most dogs on antibiotics having diarrhea (11/15) and over half of dogs with diarrhea being on antibiotics (11/21). Among NPS control dogs with clinical data available (n = 118 dogs) (**Fig. 3A**), those with diarrhea had significantly different Bray-Curtis beta diversity (p<0.001, 2.9% of total variation) and weighted UniFrac beta diversity (p<0.01, 3.7% of total variation) compared to asymptomatic animals; and those receiving antibiotics had significantly different Bray-Curtis beta diversity (p<0.001, 3.5% of total variation) and weighted UniFrac beta diversity (p<0.01, 3.9% of total variation) compared to those not receiving antibiotics, when controlling for age, sex, and spay/neuter status (**Fig. 3B**). Next, we used our NPS control group (n = 118) to define a microbiome signature associated with diarrhea in the absence of observable parasites, allowing us to compare this signature with *Giardia*-infected animals. LEfSe analysis identified *Escherichia* as enriched in animals with diarrhea and in those receiving antibiotics, while *Bacteroides* and *Fusobacterium* were enriched in asymptomatic dogs (**Figs. 3C and 3D**) and those not receiving antibiotics (**Fig S1B**). Taken together, these data define a microbiome profile associated with diarrhea and antibiotic use in NPS animals that is marked by enrichment of *Escherichia* and *Fusobacterium,* and show that this signature is distinct from that seen during *Giardia* infection (**Figs. 2B and 2C**).

**Figure 3.**
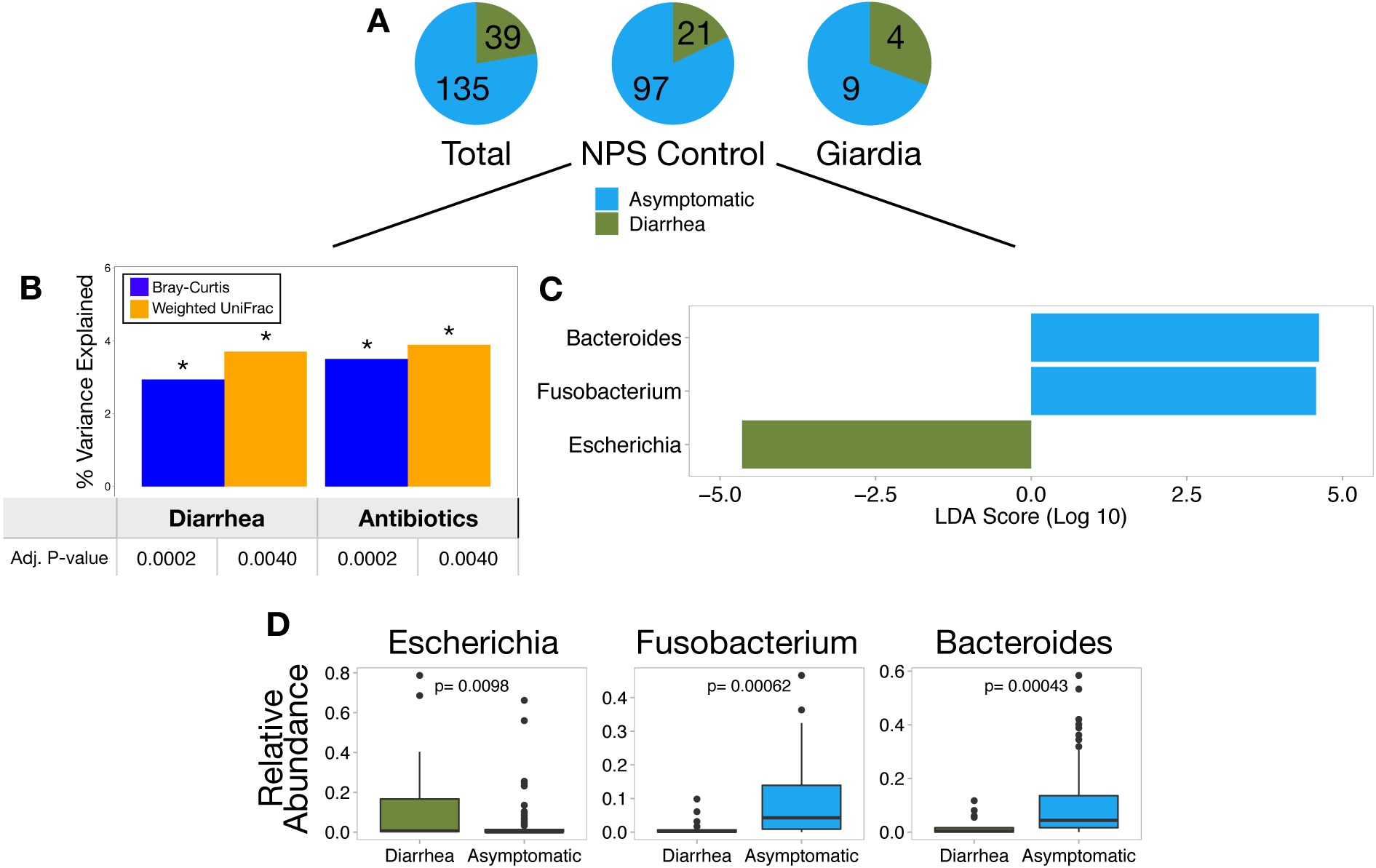
Diarrhea is associated with enrichment of a different set of taxa compared to *Giardia.* A) Pie charts show the relative proportion of sick to asymptomatic dogs among NPS Control and *Giardia*-infected dogs. The proportion of symptomatic dogs is not significantly different between groups (chi-squared p>0.05). B) Histogram shows that diarrhea and antibiotics use is associated with a significant difference in beta diversity compared to asymptomatic and no antibiotic use. Bar height reflects the percent of total beta diversity variance is explained by each variable. Age, sex, and spay/neuter status were controlled for all calculations. Asterisks denote bars with adjusted p < 0.05. C) LEfSe graph shows the magnitude of enrichment for each taxa with LDA Score > 2 comparing dogs with and without diarrhea. D) Boxplots show the relative abundance of differentially-enriched taxa. *Escherichia* is not surprisingly the most highly enriched bacterial taxa associated with diarrhea, while diarrhea is also associated with a reduction in *Bacteroides*.

### The effect of *Giardia* on the microbiome persists during co-infection

We reasoned that if *Giardia* – compared to other parasites observed in our samples – is driving changes in the microbiome, then we should observe a similar profile in animals harboring co-infections with *Giardia* and at least one other parasite. Ten out of 21 dogs harboring multiple parasites (‘co-infection’) were infected with *Giardia* and one or more other parasites. These ten *Giardia* co-infected samples were indistinguishable from *Giardia* singly-infected animals by Bray-Curtis (p>0.1) and weighted UniFrac (p>0.1) beta diversity (**Fig. 4**). In contrast, *Giardia* singly-infected samples were significantly different from the remaining 11 co-infected samples not involving *Giardia* by Bray-Curtis (p<0.05) and weighted UniFrac (p<0.05) beta diversity, however false discovery rate correction raises these p-values slightly above the 0.05 significance threshold (**Fig. 4**). Taken together, these results show that *Giardia* infection in dogs is associated with a unique and significant change in gut microbiota composition compared to NPS controls that persists even in the context of co-infection with other parasites.

**Figure 4.**
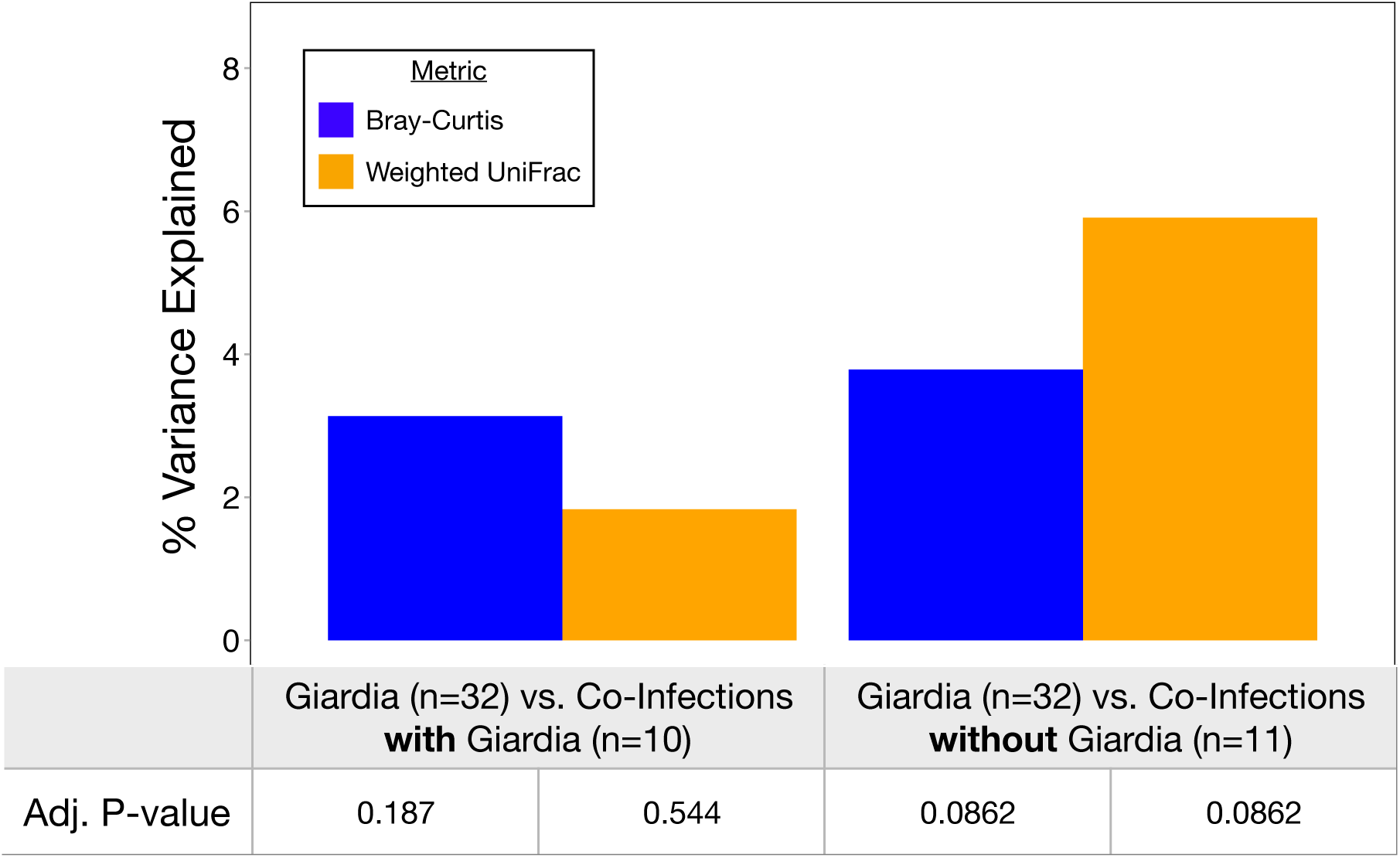
The effects of *Giardia* in canines persist even when one or more other parasites are present. Fecal samples from dogs infected with multiple parasites, one of which is *Giardia*, are not different from those singly-infected with *Giardia* in terms of Bray-Curtis or weighted UniFrac beta diversity (p>0.1). In contrast, fecal samples from dogs infected with multiple, non-*Giardia* parasites are different from those singly-infected with *Giardia* (p<0.05; adjusted p < 0.1), and infection status here represents a larger percentage of the variance in beta diversity. Age, sex, and spay/neuter status were controlled for all calculations.

### *Giardia* infection is among the largest predictors of the pediatric gut microbiota structure

After finding that parasites, in particular *Giardia*, perturb the canine gut microbiome, we asked if *Giardia* similarly affected the human gut microbiome. To this end, we employed a database mining approach to integrate and query data from the Global Enteric Multicenter Study (GEMS). Clinical and epidemiological data from GEMS was made available on ClinEpiDB.org (**Figs. 5A and 5B**) from over 22,000 participants. These data were manually combined with fecal microbiome data from a subset of the same participants (n=1004), 215 of whom were positive for *Giardia*, which was loaded on MicrobiomeDB.org (29, 30). Not surprisingly, age and moderate-to-severe diarrhea (MSD) were strongly correlated with Bray-Curtis beta diversity (p<0.001), explaining 11% and 5.5% of the total variation in microbiome structure, respectively (**Fig. 5C**). *Giardia* infection was associated with a similarly large perturbation of the gut microbiota (p<0.001; 1.9% of the total variation), while *Cryptosporidium* and Rotavirus infection were each associated with <0.5% of the variation in microbiota composition in this cohort (p<0.01).

**Figure 5.**
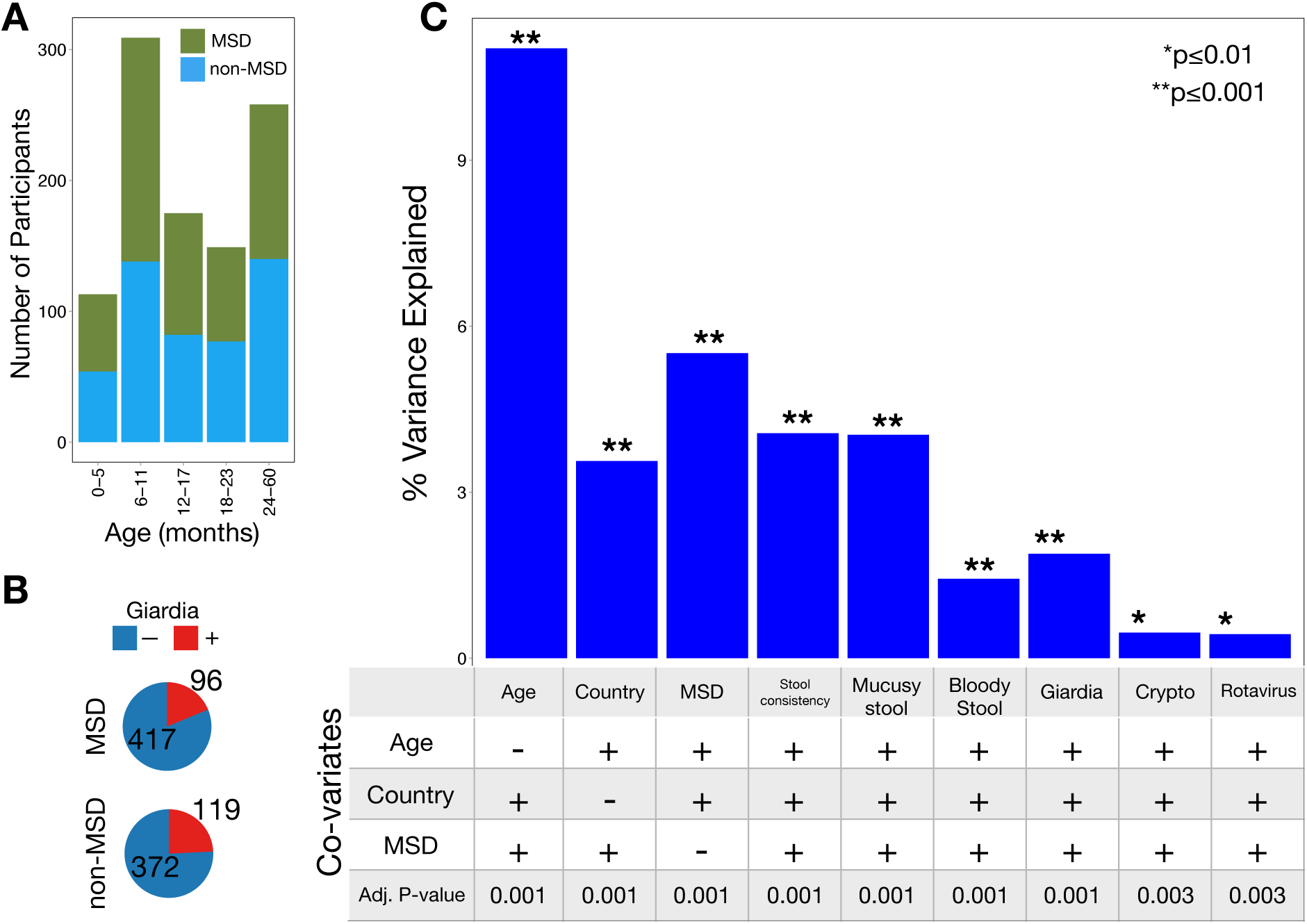
Enteric parasites are associated with gut microbiota perturbations in children. A) The number of participants with (green) and without (blue) moderate-to-severe diarrhea (MSD) in each of five age cohorts is shown B) *Giardia* is more frequently found in children without MSD compared to children with MSD. C) The percent variation in Bray-Curtis beta diversity explained by several variables is shown as bars. Whether the analysis was stratified by age, country, and/or MSD status is shown below each bar. *Giardia* is significantly associated with a change in gut microbiota, and explains more microbiota variation than any other enteric parasites and pathogens detected here.

We observed that *Giardia* infection among GEMS participants was associated with enrichment of *Prevotella* and a reduction in *Gammaproteobacteria* (**Fig. 6A**) – an effect that was evident in children with (**Fig. 6B**) and without MSD (**Fig. 6C**). Diarrhea is commonly associated with a reduction in *Prevotella* and an increased abundance of *Gammaproteobacteria*. Moreover, age strongly influences the relative abundance of *Prevotella* and *Gammaproteobacteria* (**Figs. S2A and S2B**, respectively), as well as *Giardia* prevalence (**Figs. S2C and S2D**). To control for these factors, the impact of *Giardia* was assessed among 12-17 month old GEMS participants, a cohort with high relative abundance of both *Prevotella* and *Gammaproteobacteria*, high prevalence of *Giardia* infections (29.1%; n=51), and for which *Giardia* prevalence is not correlated with age (chi-squared; p=0.99). Among 12-17 month old children, the association between *Giardia* infection and reduction in *Gammaproteobacteria* and enrichment of *Prevotella* remained (**Fig. S3**). Taken together, these results demonstrate that *Giardia* infection is associated with altered gut microbiome structure in humans and animals, marked by changes in the relative abundance of taxa linked to gut health (31–33).

**Figure 6.**
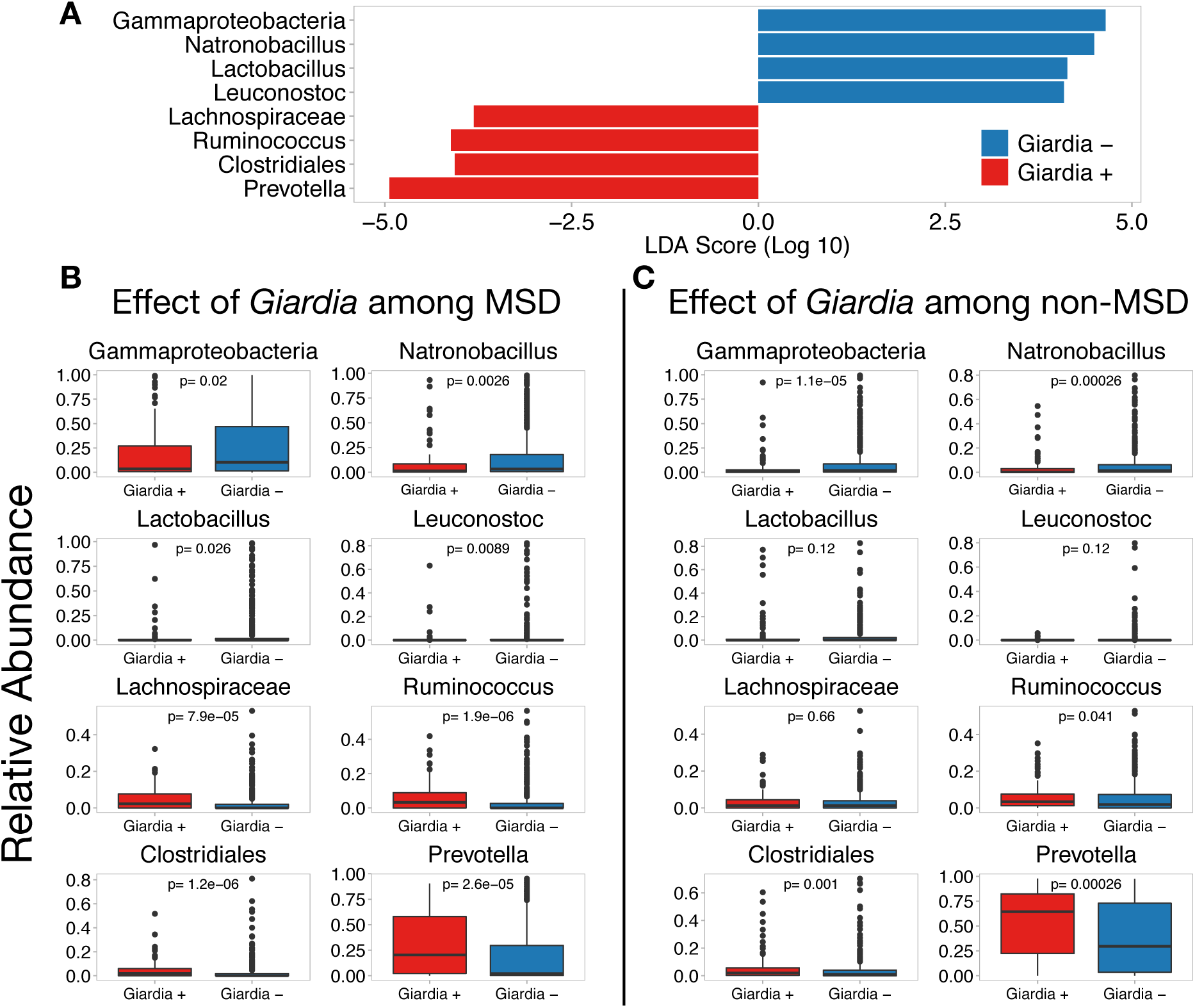
*Giardia* infection in children is associated with a reduction in *Gammaproteobacteria* regardless of disease status. A) LEfSe graph shows the magnitude of enrichment for each taxa with LDA Score > 2 comparing children with and without *Giardia* infection. B) Boxplots show the relative abundance of differentially-enriched taxa among children with MSD, C) and those without MSD. A very similar set of taxa are differentially expressed during *Giardia* infection regardless of clinical disease. Although the relative abundance of *Gammaproteobacteria* and *Prevotella* are different between MSD and non-MSD, *Giardia* infection is significantly associated with a reduction of *Gammaproteobacteria* and enrichment of *Prevotella* in regardless of MSD status.

## Discussion

Enteric parasite infections are among the most common causes of diarrhea in humans in the developing world. While bacterial infections and the gut microbiome have been well-studied, the impact of enteric eukaryotic parasites on the microbiome is not well understood, with some reports showing altered microbiome composition (14, 34–37) while others showed either modest or no impact (38, 39). Because these studies often rely on experimental infection with one or few parasite species, they provide limited insight into the broader impact of enteric parasites on the gut microbiome. By combining clinical parasitology and microbiome profiling from humans and canines infected with a phylogenetically diverse range of enteric parasites, we show that naturally-acquired enteric parasite infections are a major factor associated with microbiome composition, that this effect is observed across host species, and that *Giardia* is associated with the largest impact among all parasites surveyed in dogs and humans.

*Giardia* is one of the most common enteric parasites in the world and is remarkable in its ability to cause an array of clinical phenotypes, ranging from asymptomatic infection to severe acute diarrheal disease to chronic gastrointestinal disease. *Giardia* is the causative agent of Giardiasis, a diarrheal illness, and is clearly implicated in serious growth stunting and long-term health consequences (40), cementing its role as a pathogen. However, our observations (**Fig. 3**), as well as other reports in humans and animals suggest that *Giardia* infection is frequently asymptomatic (8, 40–47). Intriguingly, several large epidemiological case-control studies recently showed higher *Giardia* prevalence in asymptomatic participants compared to those with moderate-to-severe diarrheal disease, revealing a possible protective role (27, 40, 45, 48–50). *Giardia* infection may confer protection against diarrhea in some individuals by modulating the immune response to other pathogens (45, 51), but we reasoned that parasite-induced perturbations in the microbiome could also be an important factor influencing gastrointestinal symptoms. Our results raise the possibility that the shift in microbiome composition during *Giardia* infection – marked by a reduction in *Gammaproteobacteria* and an increase in *Prevotella* – may explain, at least in part, the apparent protective effect of *Giardia* against diarrhea in some age/site cohorts (27, 45, 50). Additional studies are needed to investigate possible consequences of interactions between *Giardia*, the microbiome, and the host. One possibility is that *Giardia* may benefit directly from manipulation of the microbiome. Interestingly, infection by another protozoan parasite, *Entamoeba histolytica,* results in enrichment of *E. coli* that protects the parasite from oxidative damage by producing malate dehydrogenase (52). Similarly, during infection with the helminth *Trichuris muris*, *Proteobacteria* directly interact with parasite eggs to induce hatching, thereby enhancing worm reproduction (53). Taken together, these studies highlight that eukaryotic parasites impact the microbiome in ways that can influence host health, immunity and parasite biology.

Our data show an association between *Giardia* infection and microbiome composition, but do not directly demonstrate that compositional changes are caused by infection. Although it may seem surprising that a pathogen of the upper small intestine could have the potential to impact microbiome composition in the stool, recent studies in mice experimentally infected with *Giardia* revealed alterations of the gut microbiome throughout the small and large intestine, indicating both a causal role and the ability to profoundly impact bacterial community structure far from the site of infection (12). The mechanisms by which this occurs have yet to be explored. Interestingly, *Giardia* infection is associated with malabsorption of fats, leading to intestinal steatosis and increased transit of lipids into the distal small intestine and colon (54), which could alter substrate availability for commensal bacteria, providing a possible explanation for compositional changes in the microbiome during this infection.

The age at which humans or animals are exposed to *Giardia* is thought to impact clinical manifestations. For example, there appears to be a window of time early in childhood development when *Giardia* infection is protective against diarrhea. Studies of several pediatric cohorts show either no correlation or a negative correlation between *Giardia* infection and diarrhea (27, 40, 42–44, 55). In contrast, adults – especially those in non-endemic areas – show a positive correlation between *Giardia* infection and diarrhea (40, 56). Previous studies suggest that the association between growth stunting and *Giardia* infection is dependent on age (57), and specifically that asymptomatic *Giardia* infection is associated with growth stunting among children older than 18 months, but not infants or in children during their first 18 months (58). The effects of *Giardia* on gut microbiota may also be age dependent. A study of Peruvian children found that gut microbiota associated with *Giardia* burden varied by age (59). For example, high *Giardia* burden was associated with enrichment of *Prevotella* only in fecal samples of 24 month old children. Here, we show an association between *Giardia* infection and altered gut microbiota composition in specific age cohorts as well, raising the possibility that parasite-microbiome interactions may partially explain the age-dependent disease presentation during *Giardia* infection. Collectively, these data point to the gut microbiome, host immunity, and age (60–62) as variables that may interact or operate independently to augment the balance between protection and pathogenesis during *Giardia* infection.

One major obstacle to investigating relationships between clinical variables and microbiome composition in large scale studies like GEMS is that these data are not always collected at the same time, by the same researchers, with the goal of being analyzed together. For example, although extensive clinical and epidemiologic data were collected from over 22,000 participants in GEMS (27), microbiome profiling data was collected from a subset of 1000 participants, and was published separately and with sparse metadata (28). Similarly, the ‘Etiology, Risk Factors and Interactions of Enteric Infections and Malnutrition and the Consequences for Child Health and Development’ study (MAL-ED) (50), also has extensive clinical metadata, along with microbiome data from a subset of participants published separately (59, 63). Our study highlights that a database-driven approach that integrates microbiome data with clinical and epidemiological data allows for the identification of novel associations and an opportunity to compare microbiome phenotypes across host species.

## Methods

### Canine sample collection

Fecal samples for our Companion Animal Microbiome during Parasitism (CAMP) study were acquired from patients seen at the Ryan Hospital at the University of Pennsylvania’s School of Veterinary Medicine (PennVet) as part of both sick and wellness visits, as well as from dogs in the spay and neuter clinic. Fecal samples were examined for parasites by fecal flotation (using a Zinc Sulfate solution at a specific gravity of 1.18) at the Clinical Parasitology Laboratory of PennVet. Dogs either had no observable parasites (n=145), one (n=92), or multiple (n=21) protozoan parasites including *Giardia* (n=32) and *Cystoisospora* (n=12), and helminths including hookworm (*Ancylostoma caninum*) (n=19), whipworm (*Trichuris vulpis*) (n=12), ascarid (*Toxocara canis*) (n=9), tapeworm (*Dipylidium caninum*) (n=5), and *Eucoleus boehmi* (n=3) (**Fig. 1A**). Samples containing yeast were excluded from the study. Age, sex, and spay and neuter status was recorded at the time of fecal sample collection for all samples. Fecal samples from 113 infected dogs and 145 dogs without detectable parasites were stored at −80C until DNA extraction. Clinical data from patient visits were obtained for 174 PennVet patients to determine whether gastrointestinal symptoms or antibiotic use occurred within one week of fecal sample collection.

### 16S rRNA gene sequencing and analysis

DNA was extracted from fecal samples using Qiagen PowerSoil DNA extraction kit. 16S rRNA sequencing was performed as described previously (64). Briefly, the V4 region of the 16S rRNA gene was amplified using PCR, which was performed using Accuprime Pfx Supermix and custom primers for 30 cycles (64). PicoGreen quantification was used to normalize post-PCR products and AMPureXP beads were used to clean the combined pools. Libraries were quantified and sized using a Qubit 2.0 and Tapestation 4200, respectively. 250bp paired-end sequencing was performed using an Illumina MiSeq. The QIIME2 pipeline (65) was used to process and analyze 16S sequencing data. Samples were demultiplexed using q2-demux and denoised using Dada2 (66). Sequences were aligned using maaft (67) and phylogenetic trees were reconstructed using fasttree (68). Weighted UniFrac (69) and Bray-Curtis (70) beta diversity metrics were estimated using q2-core-metrics-diversity after samples were rarefied to 4100 reads per sample, and p-values were adjusted for multiple hypothesis testing using Benjamini-Hochberg (B-H) false discovery rate (FDR) corrections (71). Taxonomy was assigned to sequences using q2-feature-classifier classify-sklearn (72) against the Greengenes 13-8 99% OTUs reference sequences (73). Taxa were collapsed to the genus level, when possible. OTUs with less than 1% average relative abundance across all samples were removed.

### Correlation analysis and differential feature selection

The correlation between variables such as parasite infection and microbiota composition was determined using PERMANOVA as implemented in the vegan package (74) in R (75). Differentially-abundant taxa were determined using LDA Effect Size (LEfSe) (76) and p-values were adjusted for multiple hypothesis testing using B-H FDR corrections in R. Boxplots and LEfSe plots were visualized using ggplot2 (77), patchwork (78), and ggthemes (79). Point-biserial correlation coefficients were calculated to identify differentially abundant taxa between *Giardia*-infected and NPS controls with 10,000 permutations using the indicspecies package in R (80), adjusting for multiple hypothesis testing using B-H FDR corrections (Table S1).

### Integration and analysis of GEMS data

The Global Enteric Multicenter Study (GEMS) investigated the causes, incidence, and impact of moderate-to-severe diarrhea in 23,567 0-59 month-old children in Asia and Africa (27). Clinical and epidemiological data, and anthropometric measurements for each participant were downloaded from ClinEpiDB.org (29, 30). The presence of *Giardia*, *Cryptosporidium*, and Rotavirus were determined using an antibody-based ELISA test on participant fecal samples. Additionally, sequencing of the V1-V2 region of the 16S rRNA gene was performed on stool samples from 1007 participants (28). Taxonomy was determined by classifying sequences against the Greengenes 99% OTUs reference sequences. Here, clinical data from 1004 GEMS participants was downloaded from ClinEpiDB.org and the relative abundances of bacterial taxa for the same 1004 participants was downloaded from MicrobiomeDB.org (81). The datasets were manually combined so that clinical and epidemiological data were matched to gut bacterial taxa abundance data.

Correlations between clinical variables (eg *Giardia* infection) and Bray-Curtis beta diversity were calculated using the vegan package in R. Patients were divided among 5 age groups (0-6 months, 6-12 months, 12-18 months, 18-24 months, and 24-59 months) to control for the effects associated with age. Here, associations with age were stratified by country, associations with country were stratified by age, and all other associations were stratified by age group and country (**Fig. 5C**), as done by Kotloff *et al.* 2013. Taxonomy was collapsed to the genus level, when possible, and taxa with mean relative abundance across all samples <1% were removed. Differentially-abundant taxa between *Giardia*-positive versus *Giardia*-negative and MSD cases versus controls were determined using LEfSe, adjusting p-values for multiple hypothesis testing using B-H FDR corrections. LEfSe plots and boxplots were visualized using ggplot2, patchwork, and ggthemes.

## Data Availability

All sequencing data analyzed here is publicly-available on the Sequence Read Archive (SRA) under the study accession number PRJNA594732.

## Acknowledgements

The authors would like to thank Lisa Mattei and Huanjia Zhang at the Children’s Hospital of Philadelphia for assistance in sequencing.

## Figures and Tables

**Supplemental Figure 1.**
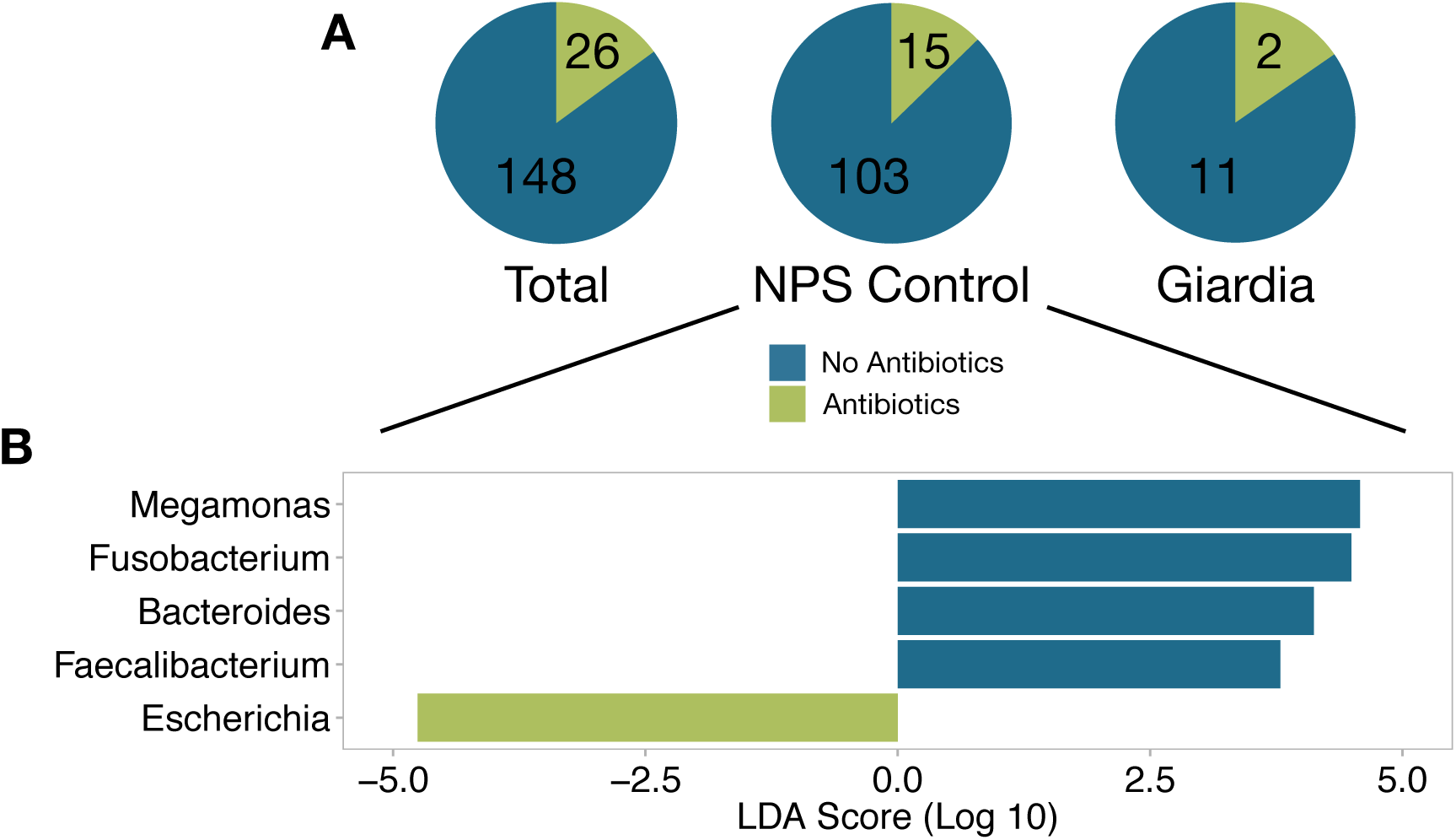
Antibiotics use in dogs is associated with a similar gut microbiota profile as dogs with diarrhea due to strong correlation between antibiotics use and diarrhea. A) Pie chart shows the relative proportion of dogs receiving antibiotics and those not among all samples that have associated clinical data (n=174), NPS controls with clinical data (n=118) and *Giardia*-positive dogs with clinical data (n=15). B) LEfSe graph shows the magnitude of enrichment for each taxa with LDA Score > 2 comparing NPS control dogs receiving and not receiving antibiotics. *Escherichia* is highly enriched in dogs receiving antibiotics and in dogs with diarrhea while *Bacteroides* and *Fusobacterium* are reduced in dogs receiving antibiotics and in dogs with diarrhea, likely because most dogs receiving antibiotics have diarrhea. *Megamonas* and *Faecalibacterium* are reduced in dogs receiving antibiotics, but not in dogs with diarrhea.

**Supplemental Figure 2.**
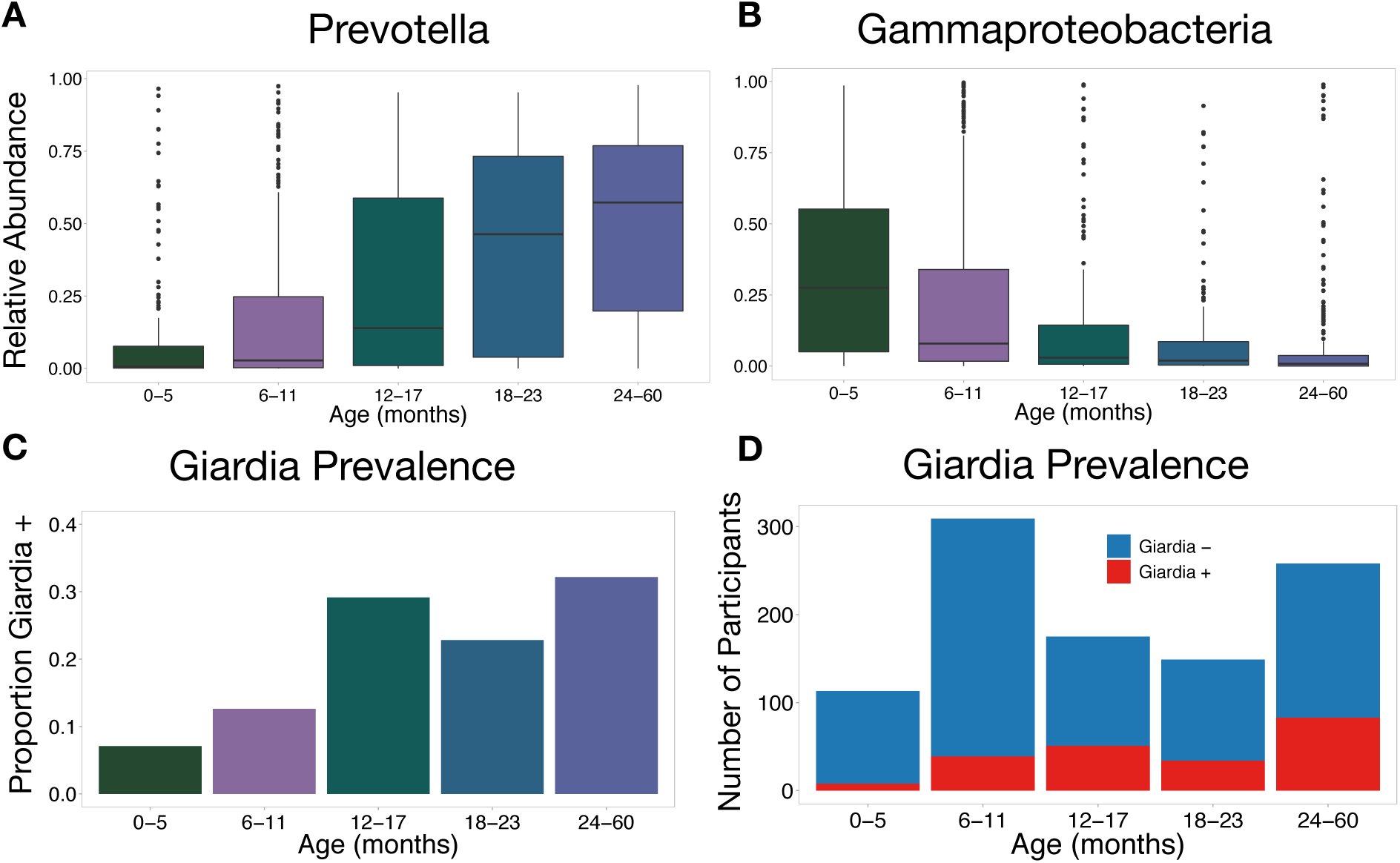
*Prevotella* abundance, *Gammaproteobacteria* abundance, and *Giardia* prevalence are correlated with age in young children. Boxplots show that, over the first 5 years of life, **A)** the relative abundance of *Prevotella* increases with age and **B)** the relative abundance of *Gammaproteobacteria* decreases with age. **C and D)** Barplots show that the proportion of children infected with *Giardia* is low for children in the first year of life compared to 1-5 year old children.

**Supplemental Figure 3.**
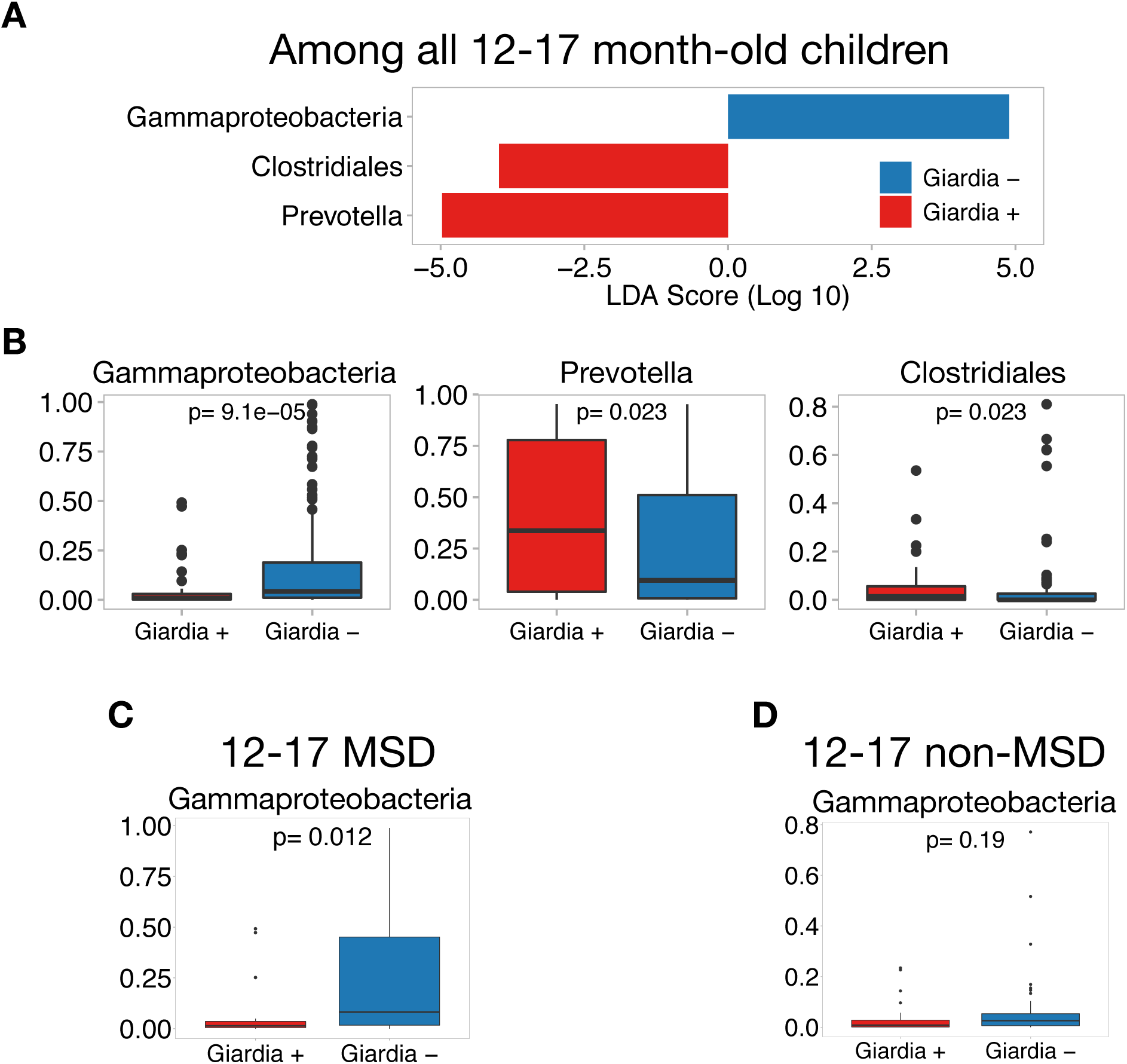
*Giardia* is associated with a reduction in *Gammaproteobacteria* and enrichment of *Prevotella* among 12-17 month old children. **A)** LEfSe graph shows the magnitude of enrichment for each taxa with LDA Score > 2 comparing 12-17 month old children with (n=51) and without (n=124) *Giardia* infection. **B)** Boxplots show the differences in relative abundance in taxa associated with *Giardia* infection among all 12-17 month old participants. **C)** Boxplots show that *Giardia* is associated with a reduction in *Gammaproteobacteria* among 12-17 month old children with MSD, **D)** but that *Giardia* is not significantly associated with *Gammaproteobacteria* among 12-17 month old children without MSD, when the relative abundance of *Gammaproteobacteria* is low.

**Table S1.** Differentially-abundant taxa associated with Giardia infection compared to NPS controls in canines identified by point-biserial correlation coefficient largely recapitulate those identified by LEfSe analysis. Point-biserial correlation coefficients show that *Clostridium*, *Lactobacillus*, and *Bacteroides* are significantly associated with *Giardia* infection, as seen in the LEfSe analysis (Fig. 2).

| **Taxa enriched in *Giardia*-infected compared to NPS Controls** |  |  |
| --- | --- | --- |
| **Taxa** | **Correlation Coefficient** | **Adj. P-value** |
| *Clostridium* | 0.29 | 0.014 |
| *Lactobacillus* | 0.23 | 0.014 |
| **Taxa enriched in NPS Controls compared to *Giardia*-infected** |  |  |
| **Taxa** | **Correlation Coefficient** | **Adj. P-value** |
| *Bacteroides* | 0.31 | 0.013 |
| *Megamonas* | 0.28 | 0.018 |

